# Age affects myosin relaxation states in skeletal muscle fibers of female but not male mice

**DOI:** 10.1101/336651

**Authors:** Lien A. Phung, Sira M. Karvinen, Brett A. Colson, David D. Thomas, Dawn A. Lowe

**Affiliations:** Department of Biochemistry, Molecular Biology, and Biophysics, University of Minnesota, Minneapolis, Minnesota, United States of America.; Department of Rehabilitation Medicine, University of Minnesota, Minneapolis, Minnesota, United States of America.; Department of Cellular and Molecular Medicine, University of Arizona, Tucson, Arizona, United States of America.

## Abstract

The recent discovery that myosin has two distinct states in relaxed muscle – disordered relaxed (DRX) and super-relaxed (SRX) – provides another factor to consider in our fundamental understanding of the aging mechanism in skeletal muscle, since myosin is thought to be a potential contributor to dynapenia. The primary goal of this study was to determine the effects of age on DRX and SRX states and to examine their sex specificity. We have used quantitative fluorescence microscopy of the fluorescent nucleotide analog 2′/3′-O-(N-methylanthraniloyl) ATP (mantATP) to measure single-nucleotide turnover kinetics of myosin in skinned skeletal muscle fibers under relaxing conditions. We examined changes in DRX and SRX in response to the natural aging process by measuring the turnover of mantATP in skinned fibers isolated from psoas muscle of adult young (3-4 months old) and aged (26-28 months old) C57BL/6 female and male mice. Fluorescence decays were fitted to a multi-exponential decay function to determine both the time constants and mole fractions of fast and slow turnover populations, and significance was analyzed by a t-test. We found that in females, both the DRX and SRX lifetimes of myosin ATP turnover at steady state were shorter in aged muscle fibers compared to young muscle fibers (p≤0.033). However, there was no significant difference in relaxation lifetime of either DRX (p=0.202) or SRX (p=0.804) between young and aged male mice. No significant effects were measured on the mole fractions (populations) of these states, as a function of sex or age (females, p=0.100; males, p=0.929). The effect of age on the order of myosin heads at rest and their ATPase function is sex specific, affecting only females. These findings provide new insight into the molecular factors and mechanisms that contribute to aging muscle dysfunction in a sex-specific manner.

## Introduction

In muscle, myosin uses energy from ATP hydrolysis to perform mechanical work with calcium (Ca^2+^) activated actin during contraction. However, when myosin is saturated with ATP in the ^2+^absence of Ca, myosin head domains are prevented from associating with actin, so the muscle remains relaxed. Recent work by Cooke and others has shown that there are two distinct states of myosin in this relaxed condition [1-3] (Fig. 1). The biochemical features of the two states were measured using a nucleotide chase experiment on skinned skeletal muscle fibers, showing that myosin single-nucleotide turnover is a multi-phasic process [1, 4]. The first relaxed state is disordered relaxed (DRX), in which myosin heads are free from each other and the thick filament backbone but do not interact with actin, resulting in orientational disorder [5, 6]. The lifetime of ATP turnover by skeletal muscle DRX myosin is less than 30 seconds (s), similar to that measured for ATPase kinetics of purified myosin [2, 7, 8]. The second relaxed state, designated super-relaxed (SRX), is proposed to be related to the highly ordered interacting head motif observed by X-ray diffraction, electron microscopy, and fluorescence polarization [3, 9-11]. In the SRX state, myosin binds ATP with extremely slow turnover lifetime (>100 s), and the two heads are proposed to be ordered in an auto-inhibitory complex. *In vitro*, relaxed fast skeletal muscle fibers typically contain ˜70% DRX and ˜30% SRX myosin [1, 2]. The SRX state is emerging as an important player in the regulation of muscle contraction and is helping address decades-old discrepancies in the myosin field. Specifically, the high rate of ATP turnover by purified myosin in solution relative to ˜10-fold slower ATP turnover in muscle fibers is explained by the decreased SRX content in purified myosin [7, 8, 12, 13].

**Fig. 1.**
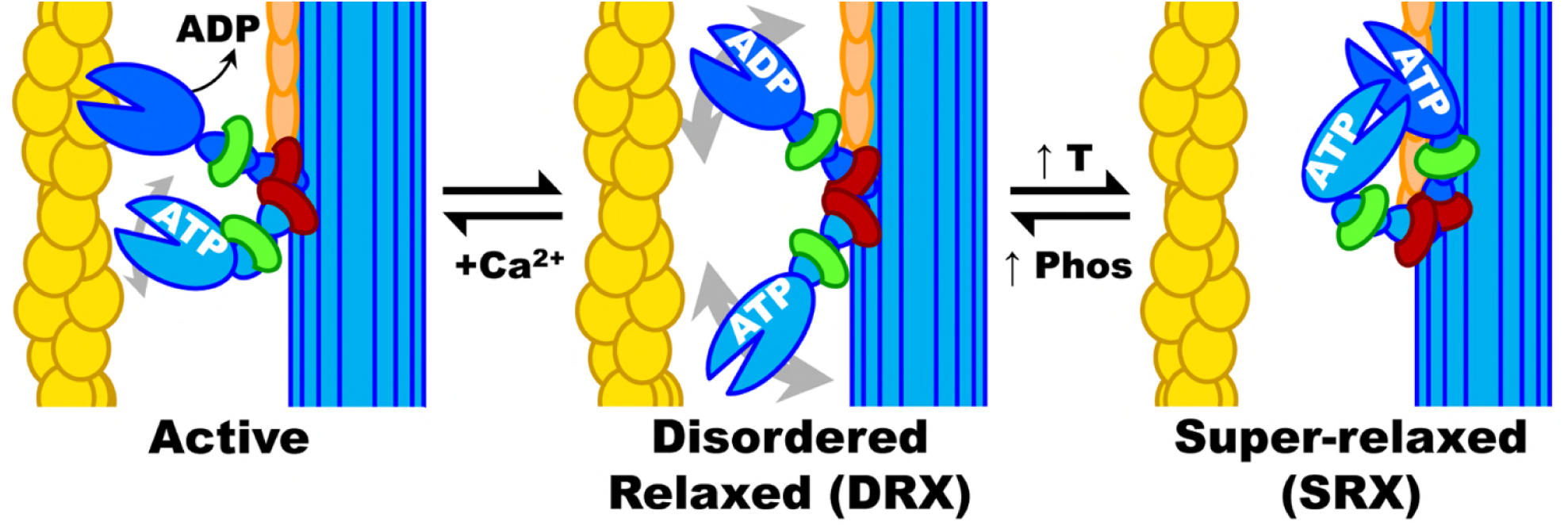
Schematic summary of myosin biochemical and structural states in muscle. Yellow: actin, blue: myosin heavy chain, green: essential light chain (ELC), red: regulatory light chain (RLC), orange: MyBP-C, Ca2+: calcium ion, T: temperature, Phos: phosphorylation of RLC and/or MyBP-C. Actin filaments are activated by Ca2+ (which binds to troponin, not shown) allowing myosin heads to be activated by actin binding followed by rapid turnover of ATP and molecular force generation. When Ca2+ is removed and ADP is replaced by ATP, myosin heads return to DRX, which can then cycle to and from SRX. When myosin heads are not ordered on the thick filament backbone (as in SRX), they are disordered and can rotate about in the interfilament space, as depicted by gray double-headed arrows in DRX.

Based on biochemical characterization of the SRX, it was observed that the myosin auto-inhibitory state can be regulated by several factors (Fig. 1). SRX can be rapidly mobilized into DRX or the active states in response to regulatory allosteric cues such as increased intracellular [Ca^2+^] or ionic strength, interaction with myosin binding protein C (MyBP-C), phosphorylation of the myosin regulatory light chain (RLC), passive strain, or decreased temperature [1-3]. Among these regulators, RLC phosphorylation is associated with muscle thermogenesis, disease, and aging [14-16].

Physiological roles of the SRX state are still under investigation, and our interest here is its possible mechanistic role in dynapenia, the age-related loss of muscle strength independent of muscle size, neurological or muscular diseases [17]. Such age-related impairments of skeletal muscle contractility are more prominent in post-menopausal women compared to men in the same age group [18]. The more rapid decline in muscle function in women has been attributed, at least in part, to the loss of ovarian hormones [18-20] and has been substantiated in studies on ovariectomized mice in which estradiol replacement improves muscle contractility [21-24]. Moreover, our group previously reported that estradiol deficiency disrupts the SRX state, supporting the hypothesis that estradiol signaling plays a role in the regulation of basal myosin function [25].

The first objective in the present study was to refine the method of analyzing fluorescence data to achieve improved measurement of the basal myosin ATP turnover activity in skeletal muscle fibers. Intrigued by how estradiol affects myosin and can reversibly regulate slow ATP turnover by SRX myosin [25], we hypothesized that muscle from aged, ovarian-senescent female mice would have altered SRX ATPase activity compared to their younger counterpart. We have tested this hypothesis using the fluorescent nucleotide chase method, and we compared male and female mice to provide insight on the relative importance of age and sex

## Methods

### Animals

Male and female C57BL/6 mice aged 3 and 22 months were acquired from the Aged Rodent Colony maintained by the National Institute on Aging (n=4 per group except aged female=7). Young adult mice were used in experiments at 3.5 to 4 months of age, and the Aged mice were kept in an SPF facility at the University of Minnesota until they reached 26-28 months of age. Mice were housed by sex in groups of 3-5 and had access to chow and water *ad libitum*. The room was maintained on a 14:10 light:dark cycle. On the day of sacrifice, mice were anesthetized by ani.p. injection of pentobarbital sodium (100 mg/kg body mass), and psoas muscles were harvested. Mice were euthanized by overdose (200 mg/kg pentobarbital). All protocols were approved by the Institutional Animal Care and Use Committee at the University of Minnesota and complied with the American Physiological Society guidelines.

### Fluorescent nucleotide exchange (“chase”) experiments

Freshly dissected mouse psoas muscles were dissected into 5 mm long bundles and chemically skinned as previously described [1, 25] and stored at −20°C for up to 3 months. On the day of experiments, single fibers were isolated from a bundle by manual dissection under relaxing conditions (4mM ATP and 4mM EGTA solution) and mounted in 35 mm glass bottom culture dishes (Bioptechs, Butler, PA) to be used with the confocal microscope. Fibers were held in place with vacuum grease placed on the ends of the fibers. Fiber segment lengths were 2-5 mm. In the present study, nucleotide exchange experiments were performed on skinned fibers with uniformly low RLC phosphorylation due to endogenous phosphatases activity. This method ensures that the results reflect the chronic state of SRX myosin without confounding effects of acute changes in RLC phosphorylation, as demonstrated in previous work [25].

All solutions in the experiment contained 120mM potassium acetate, 5mM potassium phosphate, 5mM magnesium acetate, 4mM EGTA, and 50mM MOPS, pH 6.8 [1]. This base solution is referred to as rigor buffer. Fibers were first incubated in rigor solution containing 250µM mantATP, a fluorescent nucleotide that has been shown to bind primarily to myosin in a muscle fiber (Fig. 2A) [1, 26]. After 5 minutes (min), this solution was exchanged for or “chased with” rigor buffer containing 4mM MgATP (unless noted otherwise in additional control experiments to determine the different components in the observed fluorescence decay). Each fiber was subjected to two sequential MgATP chases after washing with excess rigor buffer between runs.

**Fig. 2.**
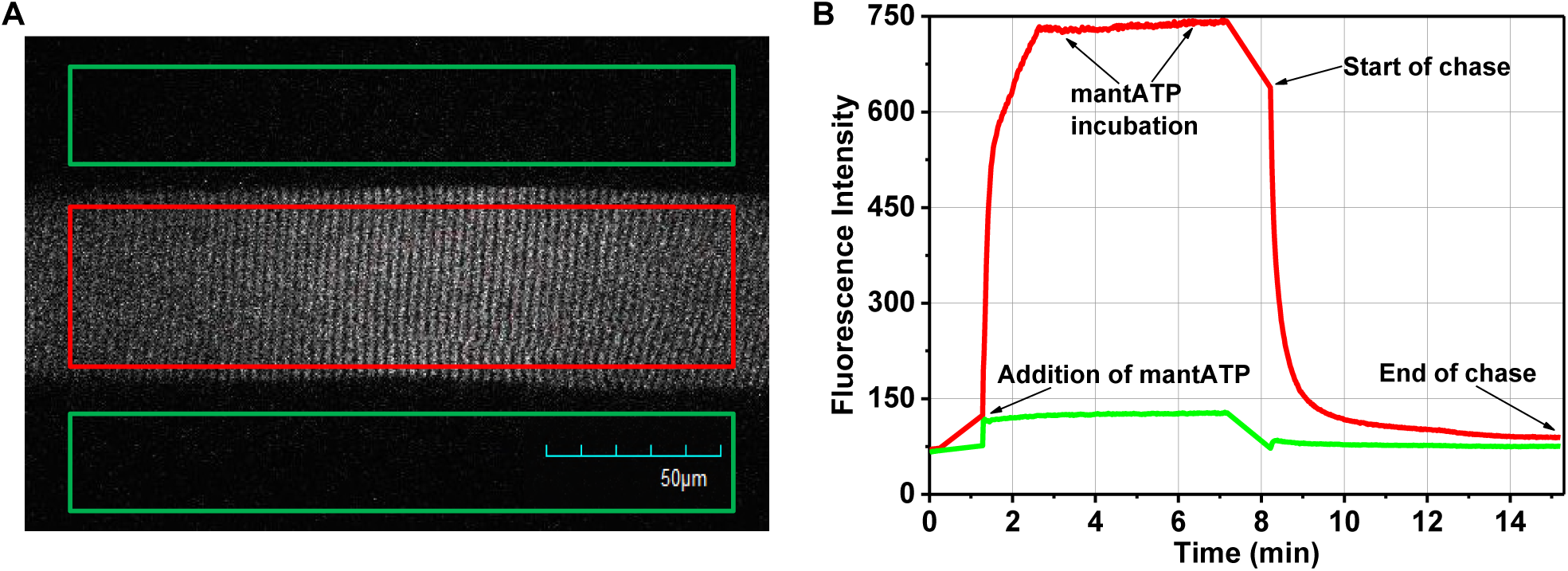
Fluorescence imaging of a muscle fiber in chase experiments. (A) Fluorescence image of a single mouse skeletal muscle fiber loaded with mantATP prior to start of ATP chase. One large rectangular region of interest (ROI) encompassing a major portion of the fiber (red) was used to monitor the time course of mantATP fluorescence intensity decay during the ATP chase. Rectangular ROI’s outside the fiber (green) were used for background subtraction, a method adapted from studies of mouse skeletal muscle fibers [25]. (B) Average fluorescence intensity of fiber (red) and background (green) ROI throughout the experiment. MantATP binds to the muscle thick filament upon addition to the culture dish. The fiber was incubated in 250µM mantATP for 5 min. Chase was started after removal of mantATP and 4mM of non-fluorescent nucleotides were exchanged in. Chase ends when fiber fluorescence intensity returns to baseline.

Experiments were performed at 21 ± 1°C with an Olympus FV1000 IX2 confocal microscope and 60x UPlanApo N oil immersion objective lens (1.42 NA) for differential interference contrast and fluorescence imaging. Temperature of the solutions were measured using a microprobe thermometer. Fluorescence was acquired with 405 nm excitation and 475/50-nm emission filters. Images were scanned in a 512 × 512 pixels grid with a total exposure time of 1.1 s and an effective pixel size of 414 nm. Fluorescence intensities of mantATP bound fiber were acquired through time-lapse imaging for 350–450 s, starting from the moment the 4mM nucleotide solution was added to the fiber. During the chase, the fluorescence intensity decreased as bound mant-nucleotides were released from myosin and replaced by non-fluorescent ATP (Fig. 2B).

Images were imported into ImageJ and average fluorescence intensities within a rectangular region that encompassed most of the fiber segment were recorded (Fig. 2A). Fluorescence signal was analyzed as the intensity in each fiber minus the average intensity outside the fiber (background) in the same image. Signals were normalized to the peak intensity value at the start of each respective chase. The data were fitted to a three-exponential decay function (Eq. 1) using a nonlinear least-squares algorithm in Origin 9.0 (OriginLab Corp.).

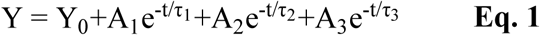

Here, A_1_, A_2_, and A_3_ are the amplitudes of each exponential term reflecting the different biochemical processes that myosin underwent during the imaging time course. These processes included fast washout of non-specifically bound nucleotides, turnover of ATP by DRX myosin, and the slow turnover of ATP by SRX myosin, respectively. τ_1_, τ_2_, and τ_3_ are the respective lifetimes of each exponential term.

Due to the close connection between muscles’ mechanical and chemical properties, even the slightest damage to the fibers’ physical structure will lead to inaccurate kinetics measurements. Therefore, each experimental fiber was subjected to careful examination for structural integrity before commencing an experiment. Also, if the macroscopic structure of the fiber changed during the course of experiment, data from that fiber was excluded from the final data set. Specifically, fibers that showed structural damage such as localized swelling, twisted or constricted areas along their length, or irregular striations were excluded. In addition, approximately 20% of the chases only fit to one exponential decay function. The data from those instances are also excluded from the final data set. The total number of fibers included in data analysis are: n=11 and 16 for young and aged female, respectively, and n= 8 and 7 for young and aged male, respectively.

### Statistical Analyses

Statistical analyses were carried out using IBM SPSS for Windows 23 statistical software (SPSS Inc., Chicago, IL, USA). The Shapiro-Wilk test was used to investigate within group normality for a given parameter of interest. Levene’s test was conducted to assess the homogeneity of the variance assumption. T-tests were used to analyze data and P-values less than 0.05 were considered statistically significant.

## Results

### Refinement of multi-exponential fit of fluorescent nucleotide signal decay

Data for the nucleotide exchange experiment were modeled with a three-exponential decay function (Fig. 3A) as described above. Previous published work on mouse and rabbit skeletal muscles used two-exponential decay fits [1, 25]. While our current data were also reasonably fitted using a two-exponential decay function, it was not the best model based on the residuals (Fig. 3B). As seen in figure 3B, a two-exponential decay function (pink) fits relative poorly in the first 120 seconds compared to a three-exponential decay function (cyan). The three-exponential function also yielded lower 𝒳^2^values by a factor of 20 on average (Fig. 3A inset). Adding a fourth exponential component did not improve the fit further (Fig. 3A, inset & 3B, purple). Thus, three-exponential fit was most appropriate for our data.

**Fig. 3.**
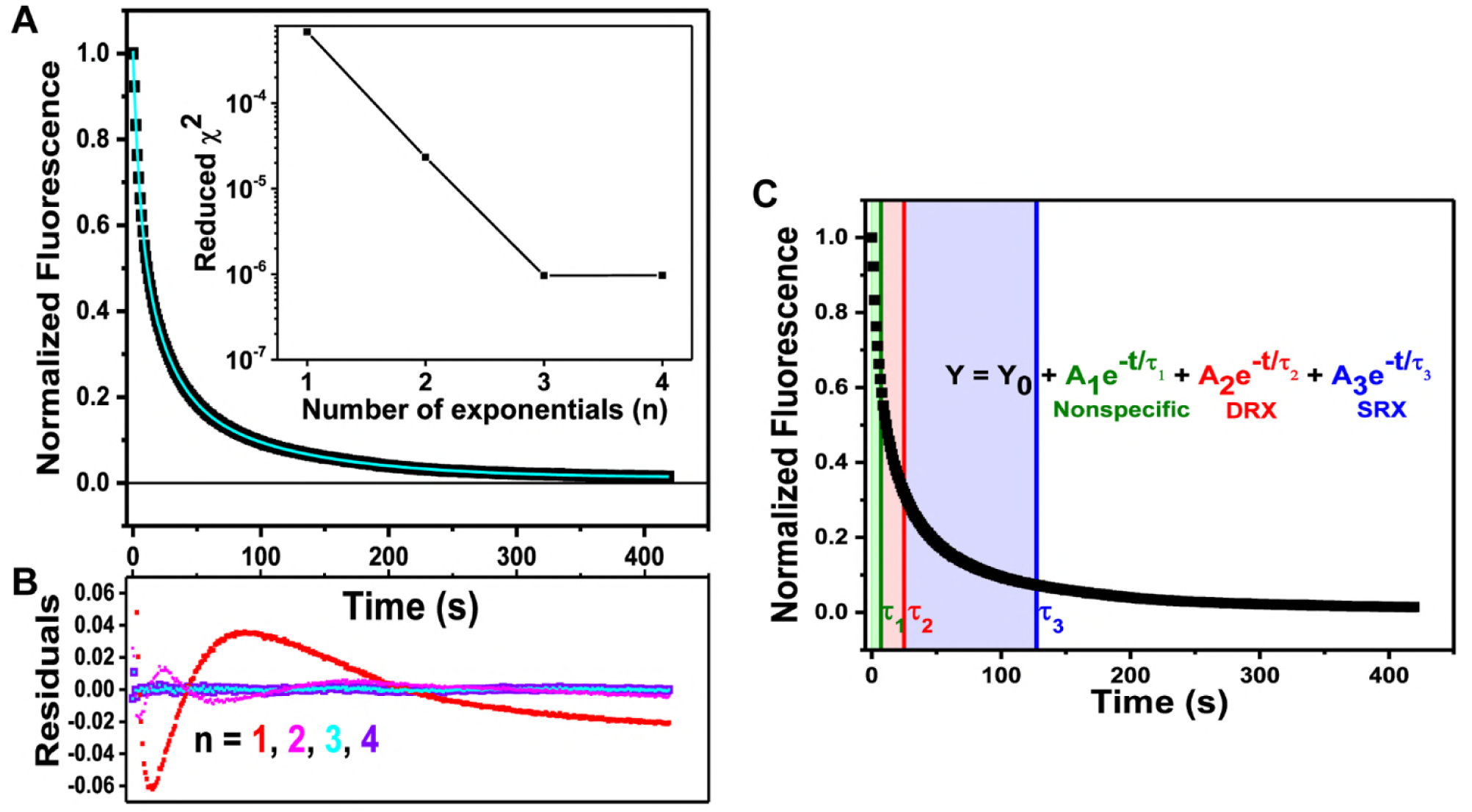
Fitting nucleotide exchange data with exponential decay functions. (A) Representative normalized fluorescence trace (black) and best fit to a 3-exponenetial decay (equation 1) from young female. Inset: 𝒳 ^2^ errors, showing that a 3-exponential fit is optimal. (B) Residual plots (fit minus data) from fits to n-exponential functions. (C) Representative trace of mantATP fluorescence decay during the time-course of nucleotide exchange with MgATP. Data from young female. All decay processes start at t=0. The color shading corresponds to the total time it takes for each kinetic process to go 63% of completion. That time is also reflected as the lifetime (τ) of each respective decay component.

The analysis method used in the current study deviates from previous studies because we used a higher sampling frequency (every second) compared to earlier works that collected data every 20 s. This affords us a better window into the nucleotide washout process that dominates the initial fast decay in the first 20 s (Fig. 3C). Lifetime of the first phase is compatible with the rapid rate of release and diffusion of nonspecifically bound nucleotides out of the fiber that occurs in ˜10 s, as previously shown [4, 14]. We postulate that the first exponential component reflects the washout of nonspecifically bound nucleotides and quick release of MgADP-P^i^from myosin heads. As an internal control, we performed sequential chase experiments where the fibers were first chased with MgATP, rewashed in rigor buffer and re-incubated in mantATP. The fiber was then chased with MgADP. After this second chase, the fiber was subjected to another round of wash and mantATP incubation for a third chase with MgATP again. Figure 4A shows representative fluorescence traces of this sequential chase experiment. The black trace is the normalized fluorescence of the first MgATP chase condition. The pink trace reflects the MgADP chase, and the blue trace is the second MgATP chase on the same fiber. The multi-exponential decay characteristic seen in MgATP chases is terminated in the MgADP chase condition, similar to previous work [1]. This results because MgADP cooperatively activates myosin and depletes the relax states [27, 28]. In MgADP chases, the myosin heads undergo rapid nucleotides release, resulting in a single-exponential decay with a lifetime of 11±2 s (Fig. 4A). Since it is possible that a small fraction of MgATP solutions contain ADP contaminants, any fast exchange of MgADP complexes will be reflected in the first phase of decay for MgATP chases. In Figure 4A, it is notable that the loss of slow myosin ATP turnover (from black to pink) can be subsequently recovered when the fiber is re-chased with MgATP (from pink to blue). This shows that myosin can undergo transitions among the different kinetic states and more importantly, that there is a population of auto inhibited, energy conserving myosin heads in relaxing conditions.

**Fig. 4.**
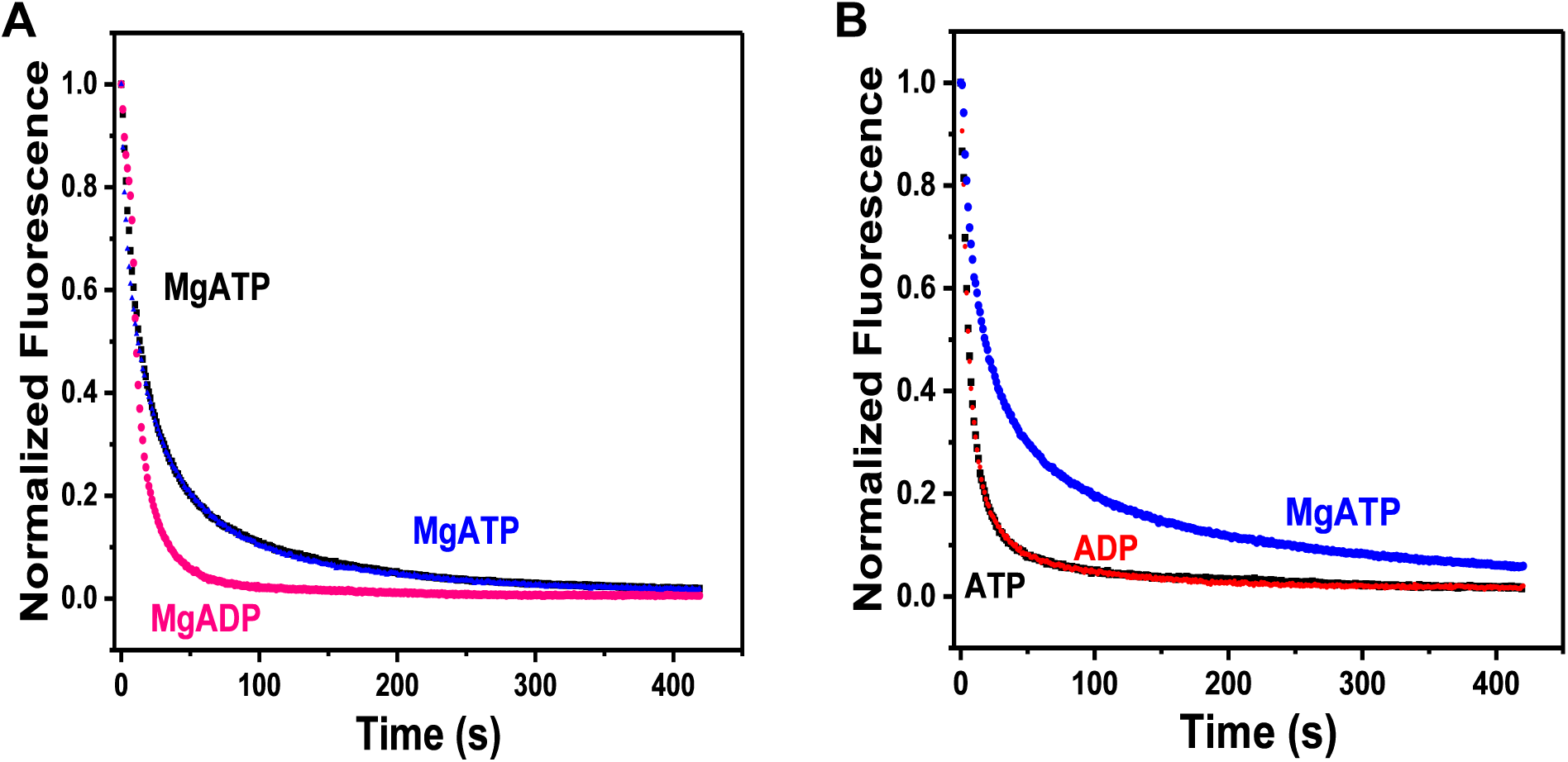
Validation of modeling nucleotide exchange with three-exponential decay function. (A) Fiber nucleotide exchange with MgATP (black and blue) or MgADP (pink). (B) Fiber nucleotide exchange in the presence (blue) and absence of magnesium using ATP (black) or ADP (red).

Given that magnesium is crucial for the coordination of specific nucleotide binding to the myosin ATPase domain [26, 29-32], we utilized this property to examine the nonspecific washout rate of nucleotides from the fibers in the chase experiments. To confirm that the first decay phase also encompasses the release and diffusion of nonspecifically bound nucleotides out of the fiber, we performed the chase experiments with and without magnesium. In the absence of the divalent ion, we expected nonspecific nucleotide binding in the skinned fibers and that the chase would exhibit a single-exponential decay with a short lifetime of less than 10 seconds. To remove magnesium from the reactions, we added 4mM EDTA to the rigor and chase buffers, and prepared fresh ATP and ADP solutions without magnesium salt. The chase experiments were carried out as described in details in the methods section above. Figure 4B shows the nucleotide exchange process on single fibers in the presence and absence of magnesium. Each tested fiber underwent sequential chases starting with 4mM ATP, then 4mM ADP, and finally 4mM MgATP. The fibers were washed in rigor buffer and re-incubated in mantATP with the respective magnesium condition between each chase. Under no magnesium conditions, the lifetime of the fastest decay was 8±2 s when chased with 4mM ATP or 7±4 s when chased with 4mM ADP (Fig. 4B). These data show that the lifetime of the fast component is independent of basal nucleotide turnover in myosin. The washout of nucleotides is a fast process that completes within the first 15 seconds of the chase experiment; thus, it is likely that the first phase would not be detected if a sampling frequency lower than 1 Hz was used.

With due diligence, we contend that the myosin nucleotide exchange in non-contracting (relaxed) mouse skeletal muscle fibers is a three-exponential process (Fig. 3C). The first component reflects the rapid release of nonspecifically bound nucleotides and their diffusion out of the fiber, with lifetime of ˜10 s. The lifetime of the second component is between 20-30 s, which is in agreement with the basal rate of myosin nucleotide turnover lifetime in solution; and therefore reflects the myosin DRX state as previously determined [1]. The third and slowest component is thus the myosin SRX state, in which turnover lifetime is generally ˜100 s. Since the first decay component is washout of nonspecifically bound nucleotides, it is appropriate to exclude the first component from the calculation of relative myosin fractions in the DRX and SRX states. Thus, fractions of DRX and SRX are derived from the amplitude values returned from the exponential decay function (Eq. 1) as detailed below. Equations 2 and 3 are fractions of DRX and SRX myosin, respectively.

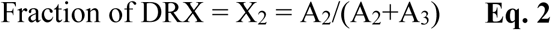

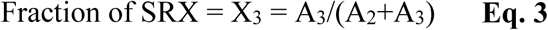

### Effect of age on SRX of skeletal muscle myosin in females

Following the previous report on the significant effect estradiol has on SRX in muscles of ovariectomized mice [25], we sought to determine if natural aging would also have negative effects on skeletal muscle myosin kinetics during relaxation. Instead of surgically removing ovaries, here mice were aged to ovarian senescence (≥26 months old) inducing a natural loss of estradiol [33]. Using young (˜4 months old) and aged (26-28 months old) female C57BL/6 mice, we tested the hypothesis that similar to ovariectomized mice, skeletal muscle myosin from aged mice would have disrupted SRX kinetics compared to young mice. In other words, we hypothesized that myosin ATPase during relaxation would be faster in fibers from aged female mice.

Applying our refined analytical approach of 3-exponential decays, we measured no effect of age on the populations of myosin in each relative state in relaxation, i.e., DRX and SRX. Here, the fraction of myosin in DRX for young and aged females is 0.74±0.01, and 0.78±0.02, respectively (p=0.100; Fig. 5A). The fraction of myosin in SRX is 0.26±0.01 and 0.22±0.02 for young and aged females, respectively (p=0.100; Fig. 5B).

**Fig. 5.**
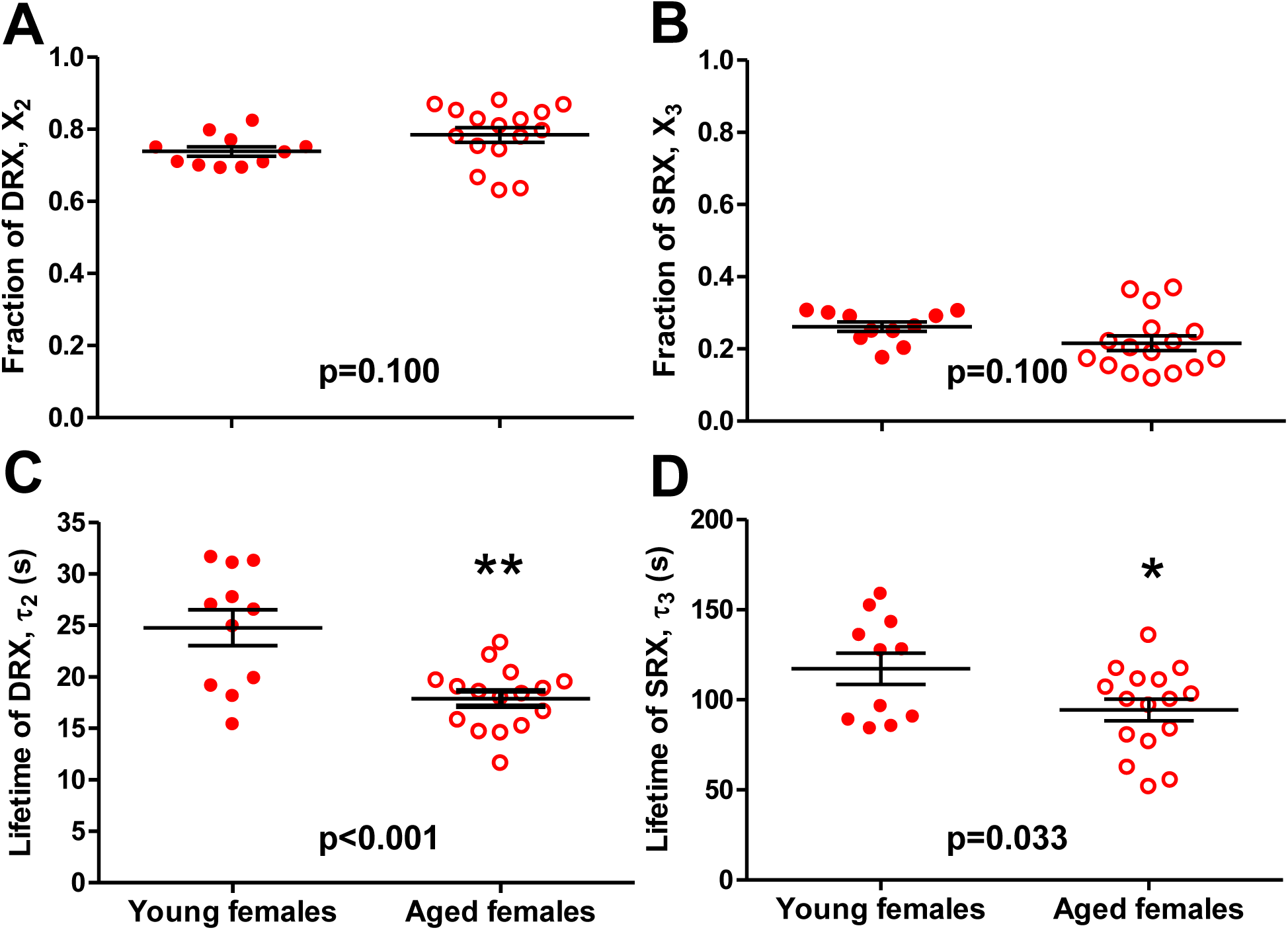
Effects of age on relaxation states of skeletal muscle myosin in fibers from female mice. The fraction of myosin (A) in DRX (X_2_ in equation 2) and (B) in SRX (X_3_ in equation 3). Lifetime of nucleotide turnover (C) in DRX (τ_2_) and (D) in SRX (τ_3_). Values are reported as average ± SEM. Individual dots are muscle fibers depicted as the average values from each chase for that fiber.

The single ATP turnover kinetics in single muscle fibers of female mice was affected by age. The lifetime of nucleotide turnover in DRX (τ_2_) was 25±2 s for young females compared to 18±1 s for aged females (p <0.001; Fig. C). The lifetime of nucleotide turnover in SRX (τ_3_) was 117±9 s for young females compared to 94±6 s for aged females (p=0.033; Fig. 5D). We conclude that myosin basal ATPase kinetics in fibers from female mice is faster with age, in both DRX and SRX.

### Effect of age on SRX of skeletal muscle myosin in males

To distinguish if the disruptive effect on myosin ATPase function is due to ovarian hormone deficiency or aging, we examined the same nucleotide turnover process in single muscle fibers from similarly aged young and old male mice. Since male mice lack ovaries and thus do not experience the same hormonal change as aging females, any observed effect on the myosin relaxed states may involve additional processes that change with age. In the male mouse fibers, we found no difference in the myosin population in each relative relaxation state between the two age groups. Here, the fraction of DRX myosin for young and aged males was 0.69±0.01 and 0.69±0.03, for young and aged males, respectively (p=0.929; Fig. 6B). Contrary to fibers from female mice, we measured no significant difference in myosin ATPase kinetics in relaxed conditions. The lifetime of nucleotide turnover in DRX was 22±1 s for young males compared to 24±1 s for aged males (p=0.202; Fig. 6C). The lifetime of nucleotide turnover in SRX was 114±7 s for young males compared to 116±7 s for aged males (p=0.804; Fig. D). In summary, there was no effect of age on myosin relaxed states in muscle fibers from male mice. This further corroborates that the disruptive effect on myosin ATPase function is related to estradiol deficiency and not specific to general aging.

**Fig. 6.**
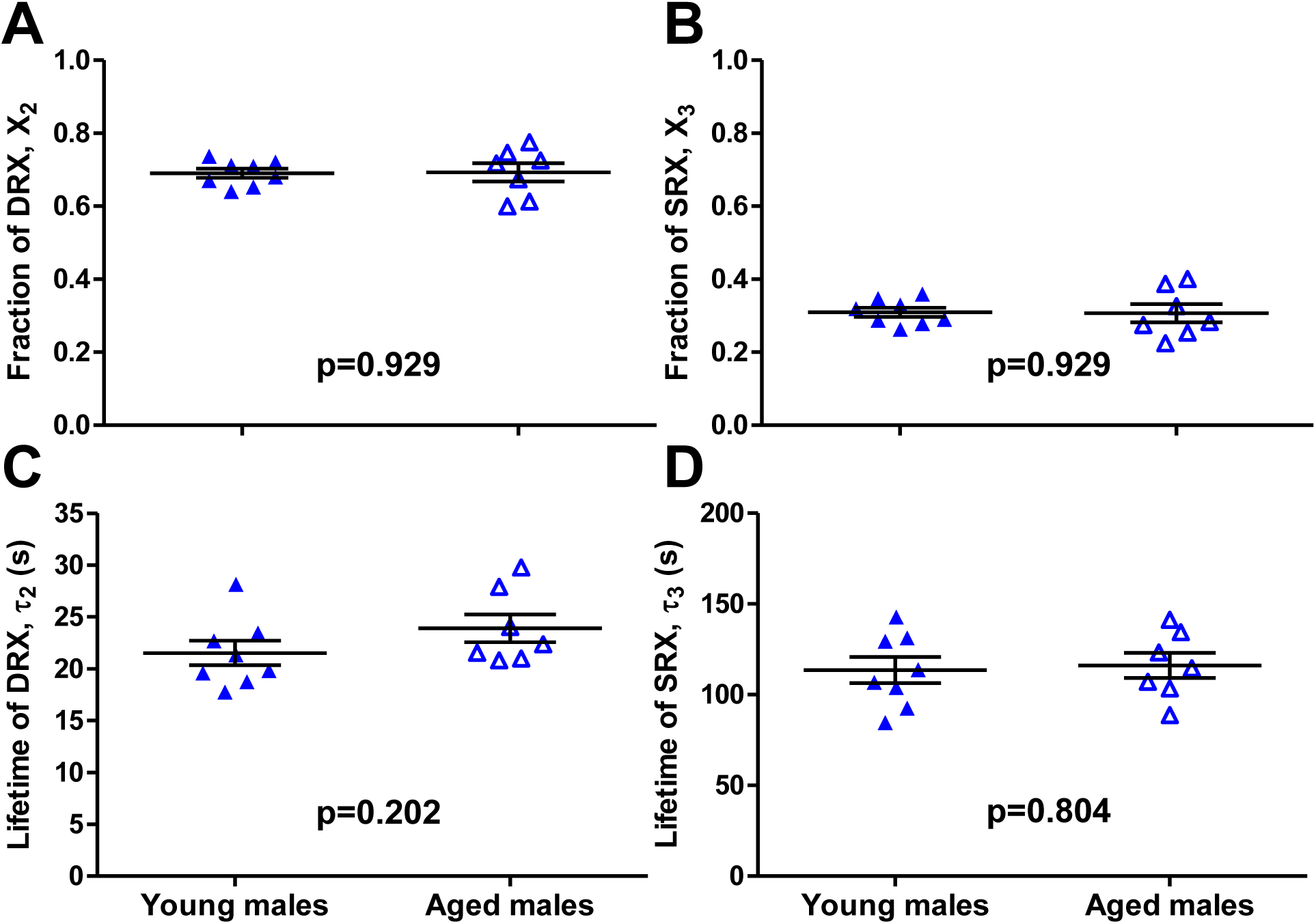
Effects of age on relaxation states of skeletal muscle myosin in fibers from male mice. The fraction of (A) DRX myosin (X_2_ in equation 2) and (B) SRX myosin (X_3_ in equation 3). Lifetime (τ) of nucleotide turnover of (C) DRX myosin (τ_2_) and (D) SRX myosin (τ_3_). Values are reported as average ± SEM. Individual dots are the average values from each tested muscle fiber.

## Discussion

This study examined the impact of the natural aging process on relaxed states of myosin in skeletal muscle fibers from male and female mice. Age-related alterations in myosin function during contraction and relaxation have been reported [15, 16, 34-39], but ours is the first to report findings for the changes in myosin ATPase activity in the DRX and SRX states with age and sex specificity. Following our previous work that showed altered SRX under relaxed conditions in fibers from an ovariectomy-induced aging model [25], here we measured the single-nucleotide turnover in resting skeletal muscle fibers isolated from naturally aged mice from both sexes. This study was done on chemically skinned fibers in fully relaxed conditions, thus the measured ATPase activity reflects mainly the work from myosin without actin activation or membrane ATPase proteins.

### Basal nucleotide turnover in relaxed skeletal muscle is a multi-component process

Because the decline in intrinsic contractile function of muscle fibers with age could be explained by age-related changes in myosin ATPase activity [16], it is pertinent to examine myosin’s basal nucleotide turnover in that context. In a minimalistic model, myosin heads in muscle fibers occupy either the active or relaxed states, the latter of which can be sub-fractionated into DRX or SRX (Fig. 1) [1-3]. Here, myosin ATPase activity in a resting muscle fiber was measured by quantitative fluorescence microscopy of mantATP turnover under relaxed condition. Applying our 3-exponential fit method, rather than the previously used 2-exponential fit [1, 25], we were able to uniquely study myosin basal ATPase activity and its specific components during relaxation (fast nonspecific washout, DRX, and SRX). Myosin in DRX turns over ATP on the order of 20-30 s in lifetime, whereas SRX myosin ATP turnover is substantially slower with an average lifetime of over 100 s. We then used the improved sensitivity to measure the kinetic parameters of DRX and SRX as they responded to the physiologic factors of age and sex.

### Basal ATPase function of myosin is altered with age in skeletal muscle fibers from females but not from males

We hypothesized that muscle fibers from aged, ovarian-senescent female mice would have disrupted SRX, represented by decreased lifetimes or fractions occupying the SRX state compared to their younger counterparts. While the fractions of SRX did not differ between young and aged female mice, we did measure a significant decrease in ATP turnover lifetime of SRX for aged female mice (Fig. 5C-D). This result agrees with our previous study where we saw a significant decrease in the lifetime of the SRX turnover from ovariectomized mice. Since slow ATPase activity is the biochemical signature of SRX, and that estradiol replacement restored the longer lifetime of the slow phase, we concluded that SRX was disrupted by estradiol deficiency [25]. In that previous study, data were analyzed using a 2-exponential decay model, so we did not have as sensitive information on DRX. The first decay phase of that method would have included the washout of nonspecifically bound nucleotides along with DRX turnover, which would have masked any measureable effect on the ATPase activity in the DRX state alone. In the present study, we used a 3-exponenital decay model and were able to measure the ATPase activity of DRX that is not confounded with the fast washout process. Interestingly, the ATP turnover lifetime in DRX was also lower in fibers from aged compared to young female mice (Fig. 5C). Together with the alteration seen in SRX, this new finding shows that the basal ATPase activity of myosin is uniformly affected by age in females. The equilibrium between the fractions of myosin in the DRX and SRX states was not affected since there was no difference in the relative myosin population in each relaxation state for young and aged mice (Fig. 5A-B).

We were also interested in the effect of age on basal myosin ATP turnover kinetics in male muscle to provide insights on the relative importance of age and sex specificity. Similar to the study in female mice, we saw no significant differences in the fractions of DRX and SRX myosin between fibers from young and aged male mice (Fig. 6A-B). The biochemical equilibrium between DRX and SRX was not perturbed by age in fibers from males either. Contrary to the effects of age seen in females, fibers from young and old male mice did not differ in the lifetime of ATP turnover in either DRX or SRX (Fig. 6C-D). Overall, there was no effect of age on myosin relaxed states or basal ATPase activity in male mice. This latter result agrees with previous studies reporting no difference in ATPase under relaxed conditions in male fibers [38, 39]. This data further corroborates that the disruptive effect on myosin ATPase function is related to estradiol deficiency and is not specific to aging process *per se*.

Estradiol-mediated effects on myosin structure and function have been investigated in both rodent models and humans. Muscle force generation and actin-myosin interactions during contraction declined in estradiol deficient mice, and estradiol replacement reversed those observed dysfunctions [20-22]. Similar results were also reported in human studies where myosin functions were altered in muscle biopsies from aged, post-menopausal women [15, 18, 19]. Mechanisms by which estradiol affects muscle fiber force-generating capacity and the dysfunctions that occur with hormone deficiency may involve post-translational modifications (PTMs) to myofibrillar proteins. Since estradiol has anti-oxidant properties [19, 40], deficiency of the hormone could accelerate oxidative stress-induced PTMs on myosin (reviewed in [41]). Qaisar et al. reported that ovarian hormone deficient women gain new PTMs as they age compared to those with hormone replacement therapy usage [19].

A PTM that is closely associated with myosin function and muscle contractility is phosphorylation, which has also been shown to be modulated by estradiol signaling [21]. Studies in rodents and women have shown that phosphorylated RLC (pRLC) decreases with age while there was no effect of age on pRLC level in males [15, 16]. We see the same trend in our current study, where muscle relaxation is affected by age in female mice, but not in males. Given that association, it is possible that RLC phosphorylation contributes to regulation of the relaxed states [1, 42]; but it is unlikely to be the main factor to explain how aged females have faster ATP turnover in both DRX and SRX. In the present study, we eliminated the acute effects of RLC phosphorylation by performing ATPase measurements on skinned fibers in which RLC phosphorylation was uniformly low, so the present effects are attributable directly to the chronic effects of aging in females. This was done to be consistent with our previous study that examined the effect of estradiol on the SRX independent of RLC phosphorylation [25].

Overall, alterations in myosin functions and structures during contraction [15] and relaxation are correlated with increased age and female sex, mirroring the demographics of dynapenic individuals [43, 44]. How the disrupted relaxation impacts the cross-bridge cycle during contraction is yet to be determined but could possibly increase the likelihood for myosin to leave relaxed states and interact with actin to produce molecular force during contraction. The subsequent actomyosin complex may stay in the strongly bound state longer, thus leading to the slowed cross-bridge kinetics and increased weakness with age measured by others [15, 16, 34-37, 45, 46].

## Conclusions

In conclusion, we have shown that the rate of myosin nucleotide turnover during relaxation is affected by age in a sex-specific manner. Specifically, age increases basal ATP turnover in muscles of female but not male mice. Together with previous results, our findings further implicate that myosin is a likely molecular contributor to dynapenia. Further studies are needed to better understand the molecular mechanisms by which myosin, its associated sarcomeric proteins, and estradiol signaling contribute to the progression of muscle weakness.

## Acknowledgements

Work was done using Olympus FluoView FV1000 IX2 Inverted Confocal Microscope and staff support at the University of Minnesota - University Imaging Centers, http://uic.umn.edu.

